# Mother–infant sharing of gut plasmid sequences during early infancy and their genetic characteristics

**DOI:** 10.1101/2024.12.19.629351

**Authors:** Yuki Kuriyama, Natsuki Suganuma, Kohei Ito

**Affiliations:** BIOTA Inc., Tokyo, 108-0074, Japan; Department of Biological Sciences, Graduate School of Science, The University of Tokyo, Tokyo 113-0033, Japan; bacterico Co., Ltd. Osaka, 530-0027, Japan; Division of Pharmaceutics, Faculty of Pharmacy, Keio University, Tokyo, 105-8512, Japan

**Keywords:** Gut microbiome, Plasmid sharing, Mother-to-infant transmission, Metagenomics, Plasmid mobilization

## Abstract

Maternal microorganisms contribute to the development of the infant gut microbiome, but plasmid sharing between mothers and infants remains poorly characterized. We reanalyzed longitudinal shotgun metagenomes from 65 mother–infant pairs to identify highly similar plasmid sequences across 117 comparisons at infant ages of 0.5, 3, and 12 months. We assessed associations with delivery mode and examined the predicted hosts and gene content of shared plasmids. Plasmid sharing was detected in 27 comparisons (23.1%) at 0.5 and 3 months, but not at 12 months. Among plasmids with predicted hosts, *Bacteroides*-associated sequences were prominent. Compared with plasmids detected only in mothers, shared plasmids were enriched in genes associated with mobilization (*mbpA, mbpB, mbpC*), replication (*repL*), and toxin–antitoxin systems (*yoeB*). Vaginally delivered infants had higher total plasmid counts at 0.5 months than cesarean-delivered infants, although plasmid sharing did not differ significantly by delivery mode. These findings provide a basis for understanding the genetic characteristics of plasmids shared during early gut microbiome development.

## Introduction

The human gut microbiome plays an important role in metabolism and immune function, and its early development may have long-term implications for human health (Afzaal et al. 2022). Maternal microorganisms contribute substantially to the initial colonization of the infant gut during and after birth (Browne et al. 2022; Vatanen et al. 2022). The composition of the early gut microbiome is influenced by several factors, including delivery mode, which affects infants’ exposure to maternal microorganisms (Mitchell et al. 2020; Dominguez-Bello et al. 2010).

Plasmids are extrachromosomal genetic elements that contribute to bacterial adaptation by carrying genes involved in functions such as antimicrobial resistance and stress tolerance (Carattoli 2013). Some plasmids can be transferred between bacterial cells through conjugation, whereas others depend on the transfer machinery of co-resident genetic elements. Bacteroidota are among the major predicted hosts of plasmids in the human gut (He et al. 2024). In addition to transfer-related functions, plasmid persistence depends on mechanisms involved in replication and maintenance, including toxin–antitoxin systems.

Maternal microorganisms acquired during early life may introduce plasmids into the infant gut, potentially contributing to subsequent genetic exchange within the developing microbial community (Pärnänen et al. 2018; Praveen et al. 2015; He et al. 2024). Previous studies have provided evidence of maternal inheritance of individual plasmids, including pBI143 (Fogarty et al. 2024). However, the prevalence and temporal patterns of plasmid sharing between mothers and infants, as well as the genetic characteristics of shared plasmids, remain poorly understood.

In this study, we reanalyzed longitudinal shotgun metagenomic data from 65 mother–infant pairs to investigate plasmid sharing during the first year of life. We examined the frequency of plasmid sharing across infant ages and its association with delivery mode. We also characterized the predicted bacterial hosts and genetic functions of shared plasmids, focusing on genes related to plasmid mobilization and maintenance. This study aims to provide an overview of mother–infant plasmid sharing and the genetic features associated with shared plasmids during early gut microbiome development.

## Materials and Methods

### Data set collection

In this study, metagenomic data (SRA: PRJNA821542) were obtained and analyzed from a previous study (Vatanen et al. 2022) that tracked gut microbiome associations between mothers and infants using longitudinal multi-omics data. Paired-end read data were downloaded using fasterq-dump (version 3.1.1) with the --split-file option. Analyses used infant samples collected at 0.5, 3 and 12 months of age together with the maternal samples collected at the matching time points; because no maternal sample was collected at 0.5 months, maternal samples collected at delivery (0 months) were used as the comparator for infants at 0.5 months. In total, 117 mother–infant comparisons from 65 pairs were available (49, 55 and 13 comparisons at 0.5, 3 and 12 months, respectively).

### Microbiome analysis of shotgun metagenomic sequences

Trim Galore pipeline (version 0.6.10) was employed for adapter trimming and quality filtering for the paired-end read data, with default parameters. To remove the human genome from quality-controlled sequences, hg38 was obtained from the UCSC genome browser (Raney et al. 2024) and mapped using bowtie2 (version 2.5.4) (Langmead and Salzberg 2012). Sequences from which the human genome was removed were assembled using Megahit (version 1.2.9) (D. Li et al. 2015) independently for each sample.

### Plasmid detection

To identify plasmids in the metagenomic sequences, SCAPP (version 0.1.1) (Pellow et al. 2021) was run with default parameters. For samples with paired-end reads out of order in forward and reverse, repair was performed with bbmap repair (version 39.01). Plasmid sequences were annotated using Bakta (version 1.9.3) (Schwengers et al. 2021) to identify protein-coding sequences (CDSs) and non-coding RNA features and assign gene names and functional annotations where available. The plasmids’ mobility type and host range were predicted using MOB-suite (version 3.1.9) with default parameters (Robertson and Nash 2018).

### Shared plasmids analysis

Plasmid sharing between mother-infant pairs was assessed using BLAST (version 2.16.0) (Altschul et al. 1990). For each family, the mother’s plasmids were used to construct a nucleotide database with makeblastdb using the options -dbtype nucl -parse_seqids. The infant’s plasmids served as query sequences in the BLAST search, which was performed with the following parameters: -perc_identity 100, -qcov_hsp_perc 90, -max_hsps 1, and -outfmt 6. A plasmid was considered shared when a single alignment covered at least 90% of the infant query sequence at 100% nucleotide identity. Persistent plasmids which those detectable at more than one age in the same infant were identified with the same procedure, using the plasmid set from 0.5 months as the database and that from 3 months as the query, for the 39 infants with assemblies at both ages.

### Statistical analyses

Plasmid counts were compared between mothers and infants, and between infants grouped by delivery mode, using the Wilcoxon rank-sum test. The association between plasmid sharing and delivery mode was assessed using Fisher’s exact test.

Annotated genetic features, including protein-coding genes and non-coding RNA features, were compared between the 34 shared plasmids and the 292 plasmids detected only in the corresponding mothers. Feature names were extracted from the gene annotation field of the Bakta output, regardless of feature type. Features without assigned names were excluded, and repeated occurrences of the same feature within a plasmid were counted once, so that each feature was scored as present or absent per plasmid. Enrichment of individual genes among shared plasmids was tested with Fisher’s exact test followed by Benjamini–Hochberg correction across the 91 annotated genetic features tested; genes with q < 0.05 were considered enriched. Odds ratios and 95% confidence intervals are conditional maximum-likelihood estimates from fisher.test. All tests were two-sided and p < 0.05 was taken as the threshold for significance. All analyses were performed in R (version 4.4.2).

## Results

### Plasmid Assembly Using Fecal Data from Mothers and Infants

We analyzed fecal metagenomic data from 65 mother–infant pairs, comprising 117 mother–infant comparisons at infant ages of 0.5, 3, and 12 months (49, 55, and 13 comparisons, respectively; Figure 1A). Plasmid contigs were recovered from both maternal and infant samples at all examined ages (Figure 1B). More plasmid contigs were generally recovered from maternal samples than from infant samples, particularly at 0.5 and 3 months (Wilcoxon rank-sum test, *p* < 0.001). At 0.5 months, infants delivered vaginally had significantly higher plasmid counts than those delivered by cesarean section (Wilcoxon rank-sum test, p = 0.046; Figure 1C).

**Figure 1.**
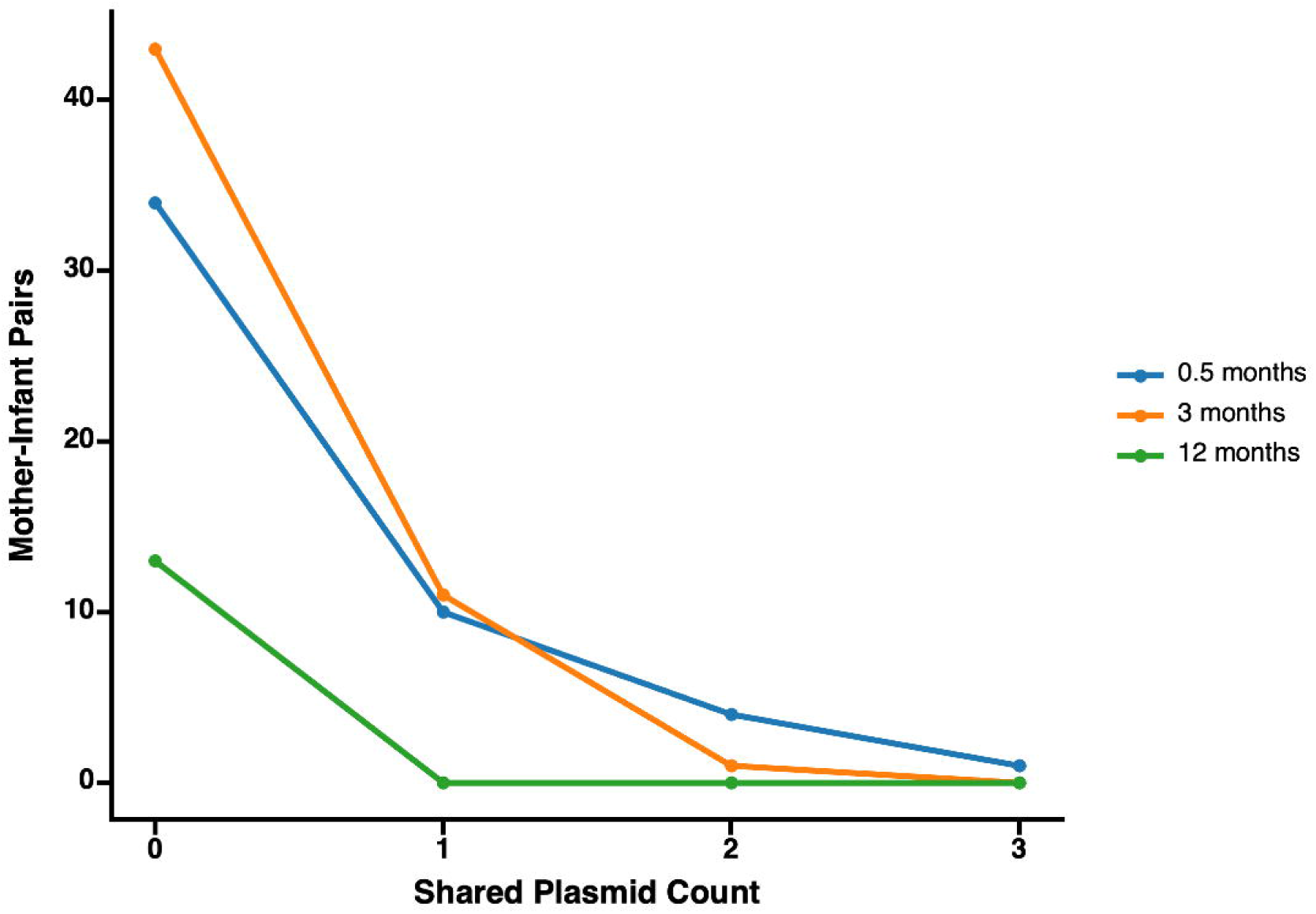
(A) Overview of metagenomic data analysis. Data from mother-infant pairs, when infants were 0.5 months (orange), 3 months (blue), and 12 months (green), were analyzed. (B) Violin plots with embedded boxplots of plasmid counts for both mothers and infants categorized by infant age. (C) Violin plots with embedded boxplots of plasmid counts for infants categorized by age and delivery mode. For all panels: *\*\*P* < 0.001; *\*P* < 0.05.

### Plasmid Sharing Between Mothers and Infants

Plasmid sharing was detected in 27 of the 117 mother–infant comparisons (23.1%), based on 100% nucleotide identity across at least 90% of the infant plasmid query sequence. The shared plasmid contigs had a mean length of 4,650 ± 2,039 bp and a mean sequencing coverage of 871 ± 1,032×. Shared plasmid sequences were detected at infant ages of 0.5 and 3 months, but not at 12 months (Figure 2).

**Figure 2.**
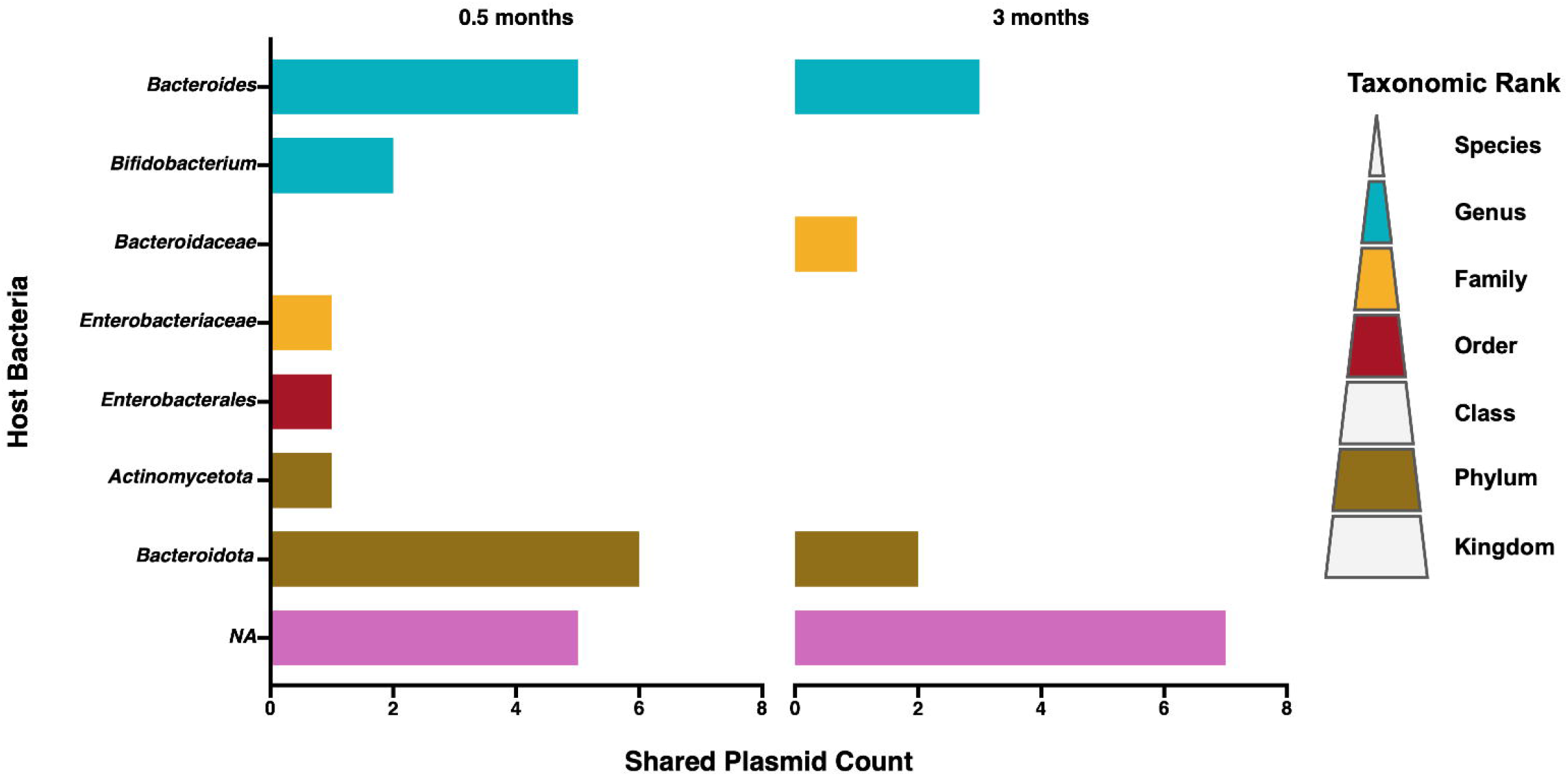
Line plots showing shared plasmid counts in mother-infant pairs across infant ages.

At 0.5 months, plasmid sharing was detected in 14 of 42 vaginal-delivery pairs and one of seven cesarean-delivery pairs (Table 1). However, the association between delivery mode and plasmid sharing was not statistically significant (Fisher’s exact test, p = 0.41). At 3 months, no sharing was detected among cesarean-delivery pairs. At 12 months, no sharing was detected in either delivery group (Figure 2).

### Host range of shared plasmids and their genetic functions

We identified 228 plasmid contigs in the 27 infant fecal samples in which mother–infant plasmid sharing was detected. Bacterial host taxonomy could be assigned to 132 contigs (57.9%). Of the 34 shared plasmids, 22 (64.7%) were assigned to bacterial taxa at varying taxonomic ranks. *Bacteroides* was the most frequently assigned genus among shared plasmids, while *Bifidobacterium*-associated plasmids were also detected at 0.5 months (Figure 3).

**Figure 3.**
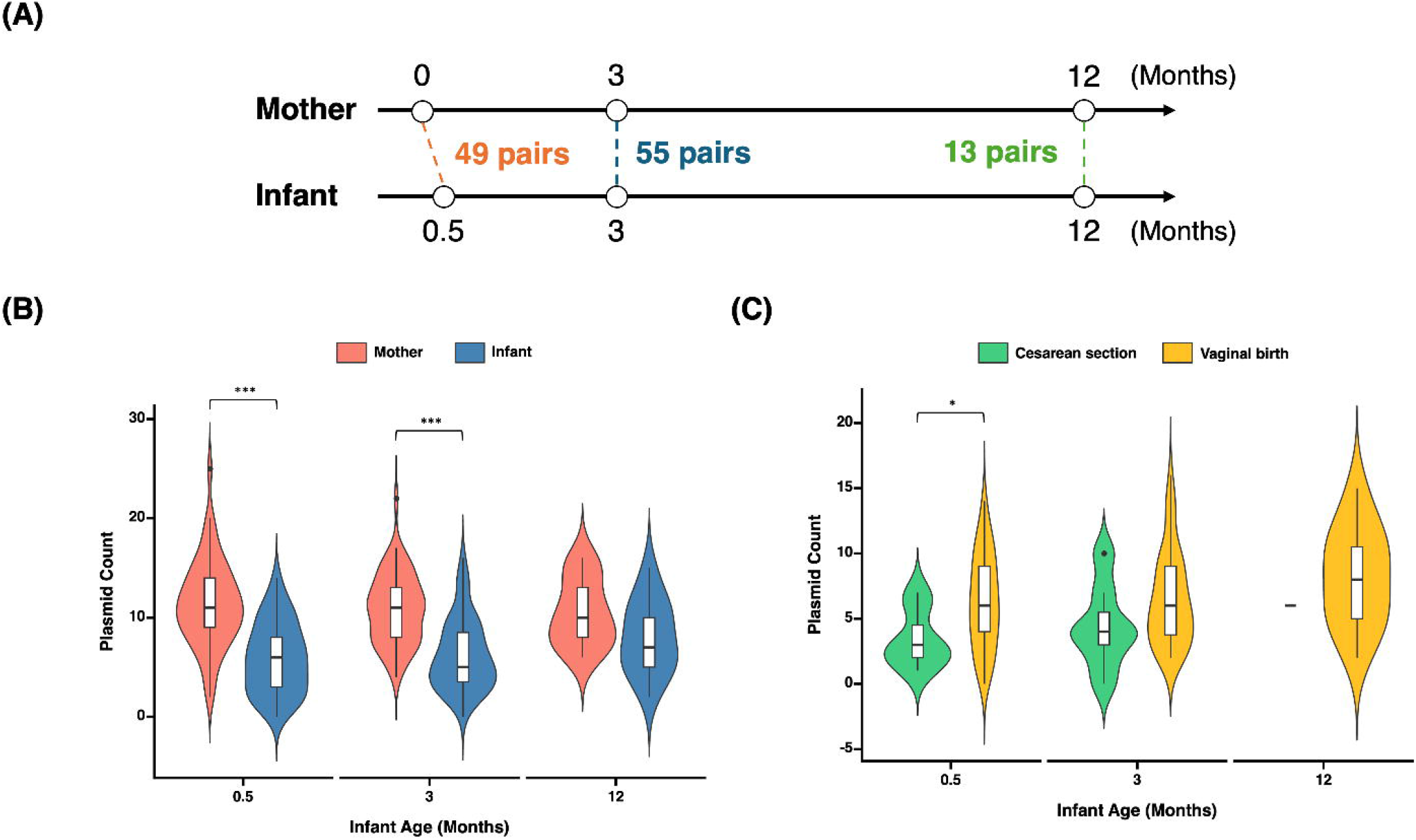
Predicted bacterial host taxonomy of plasmids shared between mothers and infants at 0.5 and 3 months of infant age. Bars indicate the number of shared plasmids assigned to each bacterial taxon, with colors representing the taxonomic rank of the assignment. NA indicates plasmids for which host taxonomy could not be assigned.

Mobilizable plasmids accounted for a higher proportion of shared plasmids (19/34, 55.9%) than unshared plasmids (85/194, 43.8%), although this difference was not statistically significant (Fisher’s exact test, p = 0.198) (Table S1). However, MOB-suite did not identify relaxases in *Bacteroides*-associated plasmids carrying the mbp mobilization cassette and classified them as non-mobilizable. When mobilization-related genes annotated by either MOB-suite or Bakta were considered, 30 of 34 shared plasmids (88.2%) carried such genes, compared with 124 of 292 plasmids detected only in the corresponding mothers (42.5%; odds ratio 10.1, 95% CI 3.4–40.5, p = 2.3 × 10^−7^). We next compared the annotated genetic features of the 34 shared plasmids with those of the

292 mother-only plasmids. Thirty shared plasmids contained at least one named genetic feature, and 91 distinct annotated features were included in the comparison. The Bacteroides mobilization genes *mbpA, mbpB*, and *mbpC* were each detected in 12 of the 34 shared plasmids (35.3%). These genes were also the most frequent among mother-only plasmids, occurring in 26, 23, and 24 of the 292 plasmids, respectively. To identify genetic features enriched among shared plasmids, we compared their frequencies between the two groups using Fisher’s exact test with Benjamini–Hochberg correction. Five genes were significantly enriched among shared plasmids: *mbpB* (OR = 6.3, 95% CI = 2.5–15.5, q = 0.002), *mbpC* (OR = 6.0, 95% CI = 2.4–14.7, q = 0.002), *mbpA* (OR = 5.5, 95% CI = 2.2–13.3, q = 0.003), *yoeB* (OR = 6.3, 95% CI = 2.3–16.4, q = 0.003), and repL (OR = 6.6, 95% CI = 2.0–20.4, q = 0.018; Table S2). No other genes were significantly enriched after multiple-testing correction.

### Persistence of Shared Plasmids in Infants

To assess plasmid persistence in infants, we compared plasmid sequences recovered at 0.5 and 3 months of age from 39 infants with data available at both time points. We identified 32 plasmids detected at both ages in 21 infants, including four plasmids shared with their mothers at 0.5 months (Figure S1). Of these four shared and persistent plasmids, three were predicted to be mobilizable and were assigned to Bacteroidota or Pseudomonadota (Table S3). Their annotated genetic functions did not differ substantially from those of the other shared plasmids.

## Discussion

In this study, we reanalyzed longitudinal metagenomic data from 65 mother–infant pairs and identified highly similar plasmid sequences in 27 of 117 mother–infant comparisons (23.1%). Plasmid sharing was detected at infant ages of 0.5 and 3 months, but not at 12 months. These findings are consistent with previous studies showing that maternal microorganisms contribute substantially to the early infant gut microbiome (Browne et al. 2022; Vatanen et al. 2022). This may also be associated with a decreasing influence of maternal gut bacteria and an increasing influence of environmental factors and diet as infants grow (Shi et al. 2024; Nielsen et al. 2020). However, the absence of detected sharing at 12 months should be interpreted cautiously because only 13 comparisons were available at this age.

At 0.5 months, infants delivered vaginally had higher total plasmid counts than those delivered by cesarean section. Although plasmid sharing was more frequently observed among vaginal-delivery pairs, the difference was not statistically significant. Previous studies have reported associations between delivery mode and early microbial colonization, including the acquisition of maternal gut bacteria (Wang et al. 2021; Ravi et al. 2018). Our findings suggest that delivery mode may be associated with the plasmid composition of the infant gut, but the small number of cesarean-delivery pairs limits further interpretation.

Plasmids contribute to the formation of the gut microbiome and are known to have diverse effects on the host by interacting with some gut bacteria (Yang et al. 2023; Zhang et al. 2022). We predicted their host range to explore the effects of shared plasmids on the gut microbiome (Figure 3). Among the plasmid contigs recovered from infants with detectable mother–infant sharing, bacterial hosts could be predicted for only 57.9% (132/228). *Bacteroides* was prominent among the predicted genera associated with shared plasmids, while *Bifidobacterium*-associated plasmids were also observed. These bacterial genera were reported in a previous study analyzing the same cohort data to be significantly more abundant in mother-to-infant strain transmission (Vatanen et al. 2022). Additionally, several extensive previous studies have reported that these genera are transmitted from mother to infant and become established over time (Browne et al. 2022; Wang et al. 2021). The large number of plasmids predicted to be hosted by *Bacteroides* and *Bifidobacterium* in the shared plasmids is consistent with these previous findings, suggesting that bacteria dominant in the gut environment may facilitate the transmission of many plasmids from mother to infant.

Plasmid mobility is an important factor in horizontal gene transfer (Smillie et al. 2010). In our study, no shared plasmids were classified as conjugative, whereas mobilizable plasmids were more frequently observed among shared plasmids (Table S1). However, some *Bacteroides*-associated plasmids carrying *mbp* mobilization genes were classified as non-mobilizable by MOB-suite, suggesting limitations in mobility prediction. Furthermore, our stringent sharing criterion of 100% nucleotide identity across at least 90% of the infant query sequence may have underestimated plasmid sharing, particularly for incompletely assembled or closely related but non-identical sequences. Larger plasmids may also have been less readily reconstructed from short-read metagenomic data, potentially contributing to the absence of conjugative plasmids among shared sequences. Notably, 86% of the reconstructed plasmids were 10 kb or smaller (median, 4,630 bp; Figure S2), indicating that our analysis primarily captured relatively small plasmids.

Among shared plasmids, *mbpA, mbpB*, and *mbpC*, which are associated with plasmid mobilization in Bacteroides, were the most frequently detected genes (Table S2). Previous experimental studies have demonstrated the involvement of these genes in plasmid mobilization (Novicki and Hecht 1995). Five genes (*mbpA, mbpB, mbpC, repL*, and *yoeB*) were significantly enriched among shared plasmids after correction for multiple testing. These genes are associated with plasmid mobilization, replication, and maintenance. However, because multiple genes may co-occur on the same plasmid, their enrichment should not be interpreted as independent functional associations. Furthermore, the predominance of *Bacteroides*-associated plasmids among shared sequences may partly contribute to the enrichment of *mbp* genes.

The enrichment of genes associated with plasmid replication and maintenance is also noteworthy. The *repL* gene is associated with plasmid replication, whereas *yoeB* encodes the toxin component of a type II toxin–antitoxin system. Such systems can contribute to plasmid stability by promoting plasmid retention in bacterial populations (Mendoza-Guido and Rojas-Jimenez 2025). These functions may be relevant during early gut microbiome development, when bacterial populations undergo substantial changes in abundance and composition. However, our analysis did not directly assess the activity of these genes or their effects on plasmid persistence. Indeed, only four plasmids shared between mothers and infants at 0.5 months remained detectable in the same infants at 3 months. Thus, although mobilization- and maintenance-related genes were enriched among shared plasmids, their contribution to plasmid transmission or persistence remains to be determined.

The enrichment of genes associated with plasmid replication and maintenance is also noteworthy. The repL gene is associated with plasmid replication, whereas yoeB encodes the toxin component of a type II toxin–antitoxin system. Such systems can contribute to plasmid stability by promoting plasmid retention in bacterial populations (Mendoza-Guido and Rojas-Jimenez 2025). These functions may be relevant during early gut microbiome development, when bacterial communities undergo substantial changes in abundance and composition. However, our analysis did not directly assess gene activity or its effects on plasmid persistence. Thus, although genes associated with mobilization, replication, and maintenance were enriched among shared plasmids, their contributions to mother–infant plasmid sharing and persistence remain uncertain.

Among 39 infants with plasmid data at both 0.5 and 3 months, 32 plasmids were detected at both ages in 21 infants. Only four of these plasmids were also shared with their mothers at 0.5 months. These observations indicate that some plasmid sequences remain detectable during early infancy, although their persistence cannot be directly linked to maternal origin. Several methodological limitations should also be considered. Our stringent sharing criterion of 100% nucleotide identity across at least 90% of the infant query sequence may underestimate plasmid sharing, particularly for divergent or incompletely assembled sequences. The use of short-read assemblies may also affect plasmid reconstruction and host prediction. Finally, without comparisons against unrelated mother–infant pairs or experimental validation, we cannot distinguish direct transmission from other explanations for sequence sharing. Thus, the findings should be regarded as descriptive evidence of plasmid sharing rather than proof of mother-to-infant plasmid transmission.

## Conclusion

In conclusion, we identified highly similar plasmid sequences shared between mothers and infants during early infancy and characterized their predicted bacterial hosts and genetic features. Shared plasmids were associated with Bacteroides and were enriched in genes related to plasmid mobilization, replication, and maintenance. Although the direction and mechanisms of plasmid transmission could not be determined, these findings provide descriptive evidence of mother–infant plasmid sharing and contribute to our understanding of plasmid diversity during early gut microbiome development.

## Acknowledgements

All authors thank Morgenrot Inc. for providing the computational environment for the analysis. We would like to express our sincere gratitude to the authors of the previous study whose data we re-analysed, particularly Dr. Vatanen and Dr. Xavier, for their generous cooperation in providing additional data.

## Authors contributions

YK collected the data set, designed the analysis workflow and performed metagenomic analysis. NS performed the genome annotation analysis and the gene-level comparison of shared and mother-only plasmids, and implemented the analysis code. KI conceived and supervised the study, designed the analysis workflow and the figures. All authors wrote the manuscript, discussed and approved the manuscript.

## Tables and figures

Table 1. Each percentage was calculated as the number of mother–infant comparisons with detected plasmid sharing divided by the total number of comparisons in the corresponding delivery-mode and age group.

Supplementary Figure 1. Venn diagram of shared and persistent plasmid counts in infants at 0.5 months. Shared plasmids were identified by comparing plasmids between mothers and their infants at 0.5 months, while persistent plasmids were determined by tracking the same plasmids in infants between 0.5 and 3 months of age.

Supplementary Figure 2. Histogram showing the size distribution of shared and unshared plasmids.

Supplementary Table 1. Distribution of predicted plasmid mobility types among shared and unshared plasmids recovered from infant fecal metagenomes. Shared plasmids (n = 34) were identified in both mothers and their infants, whereas unshared plasmids (n = 194) were detected only in infants.

Supplementary Table 2. Comparison of annotated gene frequencies between plasmids shared by mothers and infants (n = 34) and plasmids detected only in the corresponding mothers (n = 292). Genes occurring more than once on the same plasmid were counted once. Odds ratios, 95% confidence intervals, and p values were calculated using Fisher’s exact test, with q values adjusted using the Benjamini–Hochberg method.

Supplementary Table 3. Details of shared and persistent plasmids in infants at 0.5 months. The sequence coverage, predicted mobility, and host range are presented for each plasmid.

