## Supplementary Figure 1 and Figure 2 for "Mother–infant sharing of gut plasmid sequences during early infancy and their genetic characteristics"

**Supplementary Figure 2**. Histogram showing the size distribution of shared and unshared plasmids.
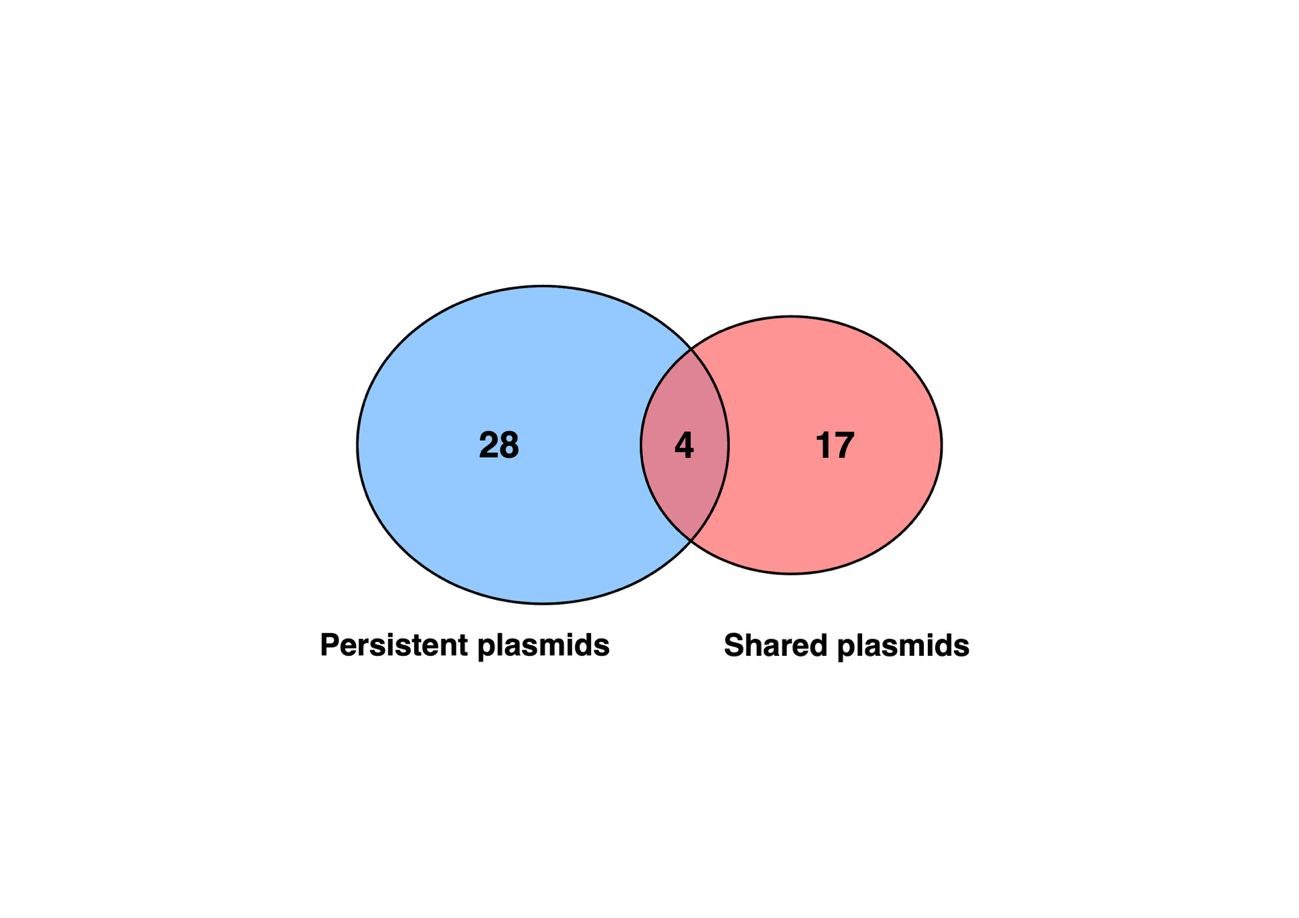

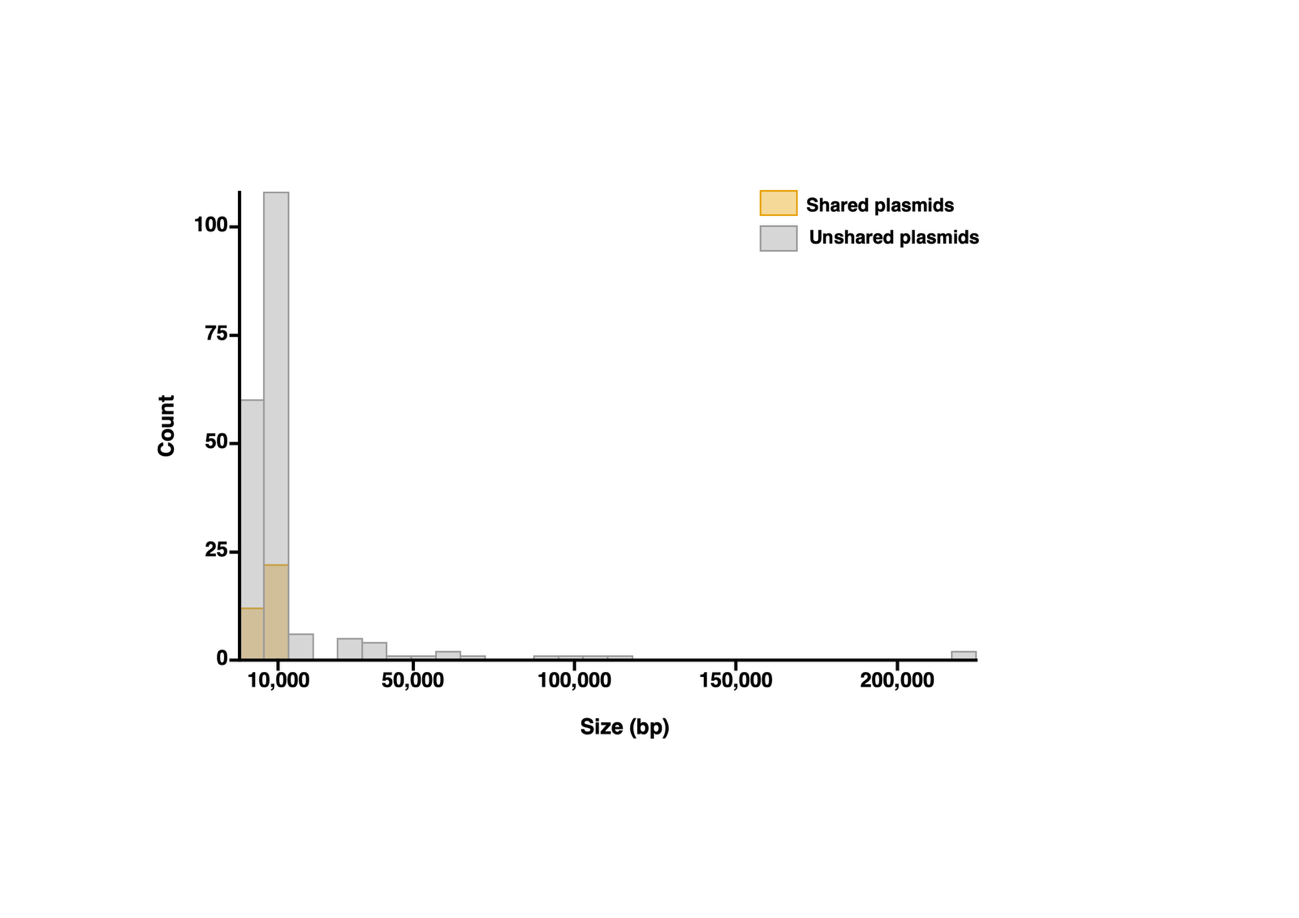

**Supplementary Figure 1**. Venn diagram of shared and persistent plasmid counts in infants at 0.5 months. Shared plasmids were identified by comparing plasmids between mothers and their infants at 0.5 months, while persistent plasmids were determined by tracking the same plasmids in infants between 0.5 and 3 months of age.
